# APRICOT: an integrated computational pipeline for the sequence-based identification and characterization of RNA-binding proteins

**DOI:** 10.1101/055178

**Authors:** Malvika Sharan, Konrad U. Förstner, Ana Eulalio, Jörg Vogel

**Affiliations:** Institute for Molecular Infection Biology, University of Würzburg, Würzburg, Germany; Core Unit Systems Medicine, University of Würzburg, Würzburg, Germany

## Abstract

RNA-binding proteins (RBPs) have been established as core components of several post-transcriptional gene regulation mechanisms. Experimental techniques such as cross-linking and co-immunoprecipitation have enabled the identification of RBPs, RNA-binding domains (RBDs), and their regulatory roles in the eukaryotic species such as human and yeast in large-scale. In contrast, our knowledge of the number and potential diversity of RBPs in bacteria is poorer due to the technical challenges associated with the existing global screening approaches.We introduce APRICOT, a computational pipeline for the sequence-based identification and characterization of proteins using RBDs known from experimental studies. The pipeline identifies functional motifs in protein sequences using Position Specific Scoring Matrices and Hidden Markov Models of the functional domains and statistically scores them based on a series of sequence-based features. Subsequently, APRICOT identifies putative RBPs and characterizes them by several biological properties. Here we demonstrate the application and adaptability of the pipeline on large-scale protein sets, including the bacterial proteome of *Escherichia coli.* APRICOT showed better performance on various datasets compared to other existing tools for the sequence-based prediction of RBPs by achieving an average sensitivity and specificity of 0.90 and 0.91 respectively. The command-line tool and its documentation are available at https://pypi.python.org/pypi/bio-apricot

## INTRODUCTION

Ribonucleoproteins and RNA-binding proteins (RBPs) are important post-transcriptional regulators in several processes such as, RNA splicing, transport, localization, translation and stabilization. Such regulatory mechanisms involve brief interactions or stable bindings of regulatory RNAs with RBPs, which are structurally and functionally important for various cellular processes. Due to developments in high-throughput mass-spectrometry and sequencing approaches, it is technically possible to perform global analyses to comprehensively catalogue RBPs in an organism. Several studies have been conducted to identify and characterize RBPs as post-transcriptional regulators in human, mouse and yeast (1–6). More than 1,000 eukaryotic RBPs have been described to contain conserved amino-acid motifs or RNA-binding domains (RBDs), which serve as RNA binding sites (1, 7). A large number of these RBDs are classified based on their RNA-binding characteristics as classical RBDs, and non-classical RBDs (1, 8) based on their identification in several RBPs or few well-characterized ribonucleoproteins respectively. Additionally, a small number of RBPs lacking known RNA-binding motifs have been identified, which in most cases rely on intrinsically disordered domains for their interaction with RNAs (1). Moreover, numerous structures of protein-RNA complexes have also been solved experimentally, providing biophysical information on the interaction between nucleic acids and amino acids.

The developments in RNA and RBP research have provided reliable resources for the advancements of computational methods for the identification of similar RBPs in different genomes. Bioinformatic approaches have been established to predict and characterize known RBPs using sequence-based features, such as biochemical properties, structural properties and their evolutionary relationship (9–11). A few computational tools such as SPOT-Seq (10), RNApred (12) and catRAPID signature (13) allow the identification of RBPs directly from the primary sequences of proteins. Other computational methods, such as RNAProB (14), BINDN+ (15) and RNABindRPLUS (16) have been developed to characterize RBPs by predicting RNA-binding residues derived from the known protein-RNA structures. Such tools can also be used to identify RBPs when RNA-binding residues in the query proteins are recognized. Since these methods are computationally expensive and have been trained on specific subsets of RBP structures, they do not perform equally well on heterogeneous datasets (17). For example, RBRIdent (18) is a recent approach that utilizes several biological features for an improved sequence-based prediction of RNA-binding residues, which, like many other tools, performs well only on specific datasets (17).

Since the experimental techniques established for the eukaryotic systems cannot be directly applied to bacterial systems without their intensive optimization, there is a lack of a system wide study of RBPs in bacteria (19). Current knowledge of the RBPs in bacterial species is restricted to only a few proteins such as Hfq and CsrA, which together with their targets are an integral part of large post-transcriptional regulons (20–24). In contrast to the limited number of RBPs in bacteria, several hundreds of non-coding RNAs have been discovered that are linked to various regulatory processes such as expression of specific regulons and transcription factors via interactions with mRNAs and proteins (20). In order to understand the mechanisms involved in such RNA-regulated events, it is crucial to quantify and characterize the proteins that interact with these regulatory RNAs. Based on the experimentally derived RBPs from all the domains of life, computational methods can be developed that are capable of screening large protein sets.

We report APRICOT, an integrated pipeline for the sequence-based identification of RBPs in complete proteome sets of both eukaryotic and bacterial species. The pipeline characterizes a protein as RBP on the basis of experimentally annotated functional motifs and domain families such as RBDs. APRICOT measures similarity between the predicted RNA-binding site in the query proteins and their corresponding reference domains based on the sequence-based features and performs statistical analyses. This tool is built upon a broad knowledge and sophisticated computational approaches in the field of functional motif discovery and our experiences of working with RNA-binding proteins in bacteria. The pipeline has been trained and tested on several test sets from protein databases and compared with previously described tools for RBP predictions. By analysing the complete proteomes of human and *Escherichia coli* (strain K-12) we demonstrate the ability of the pipeline to process large datasets including bacterial proteomes. Additionally, by easily adapting the pipeline for the identification of kinases, we demonstrate its application in the characterization of proteins by other functional classes as well.

## MATERIAL AND METHODS

### Databases and the tools

APRICOT requires a set of query proteins as input for which the presence of RBDs should be determined. The basic information, for example, amino acid sequences and taxonomy data, are retrieved from UniProt Knowledgebase (25). In addition, a reference domain set is collected from domain databases based on functional classes specified by the users.

The domain resources used in this study are, Conserved Domain Database (26) (CDD) and InterPro (27), which consist of predictive models and signatures representing protein domains, families and functional sites from multiple publically available databases. CDD includes domain entries as Position-Specific Score Matrices (PSSM) that are generated from multiple sequence alignment (MSA) of representative amino-acid sequences obtained from several domain databases, namely Pfam (28, 29), TIGRFAM (30), SMART (31), COGs (32), several NCBI curated domains like PRK or Protein Clusters (33) and multi-model superfamilies of proteins (26). For the identification of domains in a given protein sequence, the PSSM entries in CDD are queried via Reverse Position-Specific BLAST (RPS-BLAST), a variant of popular Position-Specific Iterative BLAST (PSI-BLAST) (33). CDD (v3.14) contains annotations for 50,648 domains where entries from every domain resource are assigned an individual PSSM identifier allowing redundant entries of domains.

InterPro is a similar consortium that consists of domain entries as predictive models and signatures obtained from different databases, namely Pfam (28, 29), TIGRFAMs (30), SMART (31), PROSITE patterns and profiles (35), HAMAP (36), PRINTS (37), PIRSF (38), ProDom (39), PANTHER (40), GENE3D (41) and SUPERFAMILY (42). Most of these databases contain domain entries as Hidden Markov Models (HMM) (43) probabilistic models derived from sequence alignments, which capture information on both substitution and indel frequencies. These domains can be queried using tools like HMMER3 (44). Few member databases contain PSSM domain models built from the multiple alignments of representative amino-acid sequences from the UniProt protein database, which can be queried by BLAST-based methods or single model search algorithm (45), which have been integrated into InterProScan 5 (45). As of May 2016, InterPro (v.57) contained 29,175 domain models of which several are annotated with Gene Ontology (GO) terms (46).

InterPro and CDD consortiums have only three databases in common (Pfam, TIGRPFAM and SMART) that account for about 20,000 domains. Technically, the PSSM based approach by CDD is built upon ungapped motifs, whereas the HMM probabilistic models of InterPro can handle motifs with insertions and deletions. By combining the predictive abilities of the CDD and InterPro consortiums, APRICOT provides a broader scope for domain characterization.

### Workflow

APRICOT involves different modules for the identification and characterization of RBPs, which can be explained by its program input, analysis modules and program output (Figure 1 Architecture of APRICOT.(A) A simplified overview of the processes involved in APRICOT analysis. (B) Flow-chart showing different components of APRICOT pipeline for the characterization of RNA-binding proteins. Modules for the primary analysis involving the processing of user-provided inputs (orange boxes) and the downstream analysis, which includes modules for the identification of RBPs candidates (grey box) and the modules for the annotation and feature-based scoring of putative RBPs (purple boxes). APRICOT generates a comprehensive results for each analysis, which are represented by means of tables and visualization files (green box).1). These modules are assembled into a command-line tool, for which the individual modules accessible through subcommands are specified below.

**Fig. 1.**
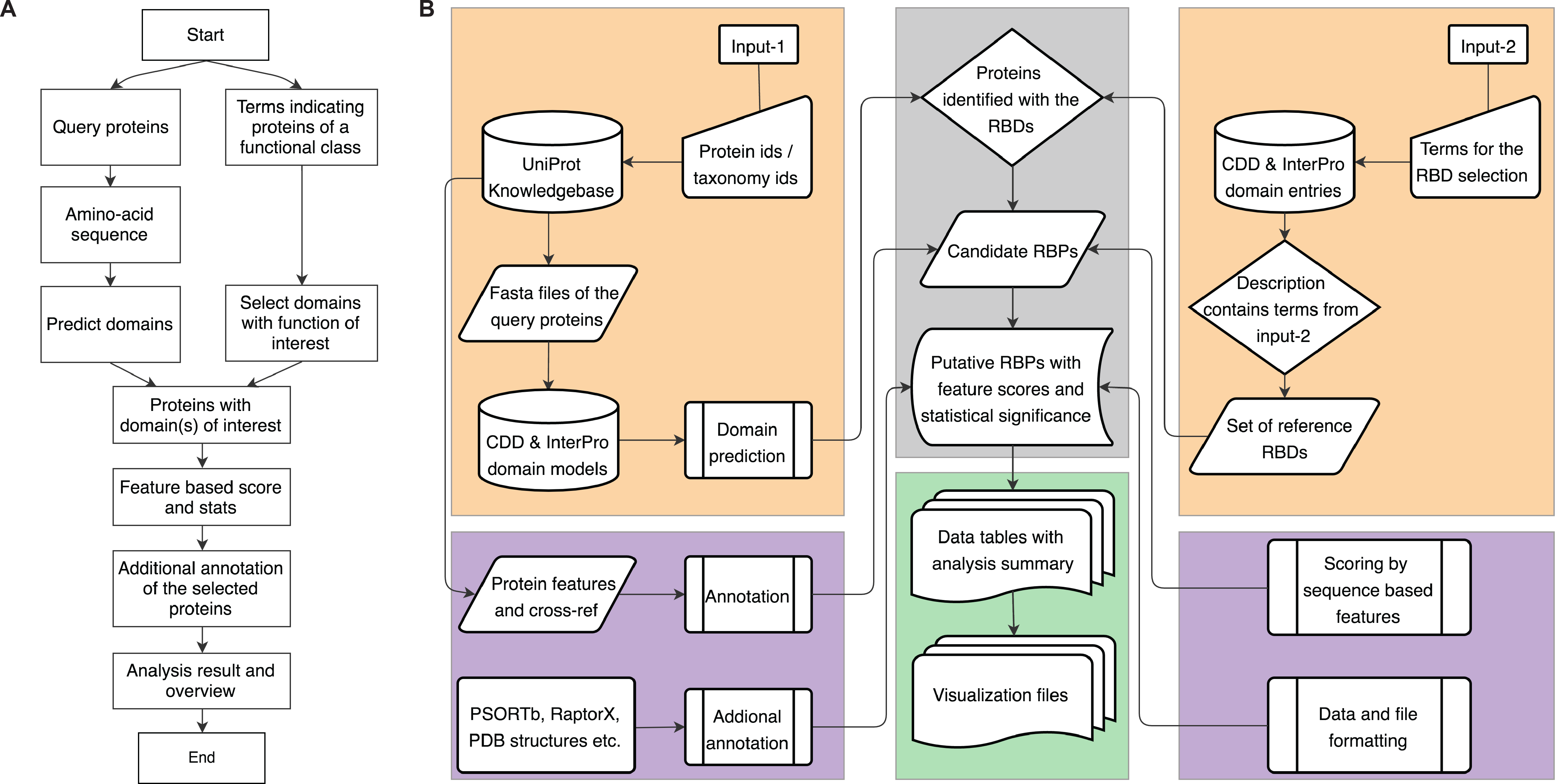
Architecture of APRICOT. (A) A simplified overview of the processes involved in APRICOT analysis. (B) Flow-chart showing different components of APRICOT pipeline for the characterization of RNA-binding proteins. Modules for the primary analysis involving the processing of user-provided inputs (orange boxes) and the downstream analysis, which includes modules for the identification of RBPs candidates (grey box) and the modules for the annotation and feature-based scoring of putative RBPs (purple boxes). APRICOT generates a comprehensive results for each analysis, which are represented by means of tables and visualization files (green box).

### 1. Program input

APRICOT requires two inputs for its execution: query proteins and the functional class of interest (Figure 1). The query proteins can be provided either as a list of gene ids, protein ids or amino acid sequences. The query search can be limited to a specific species by providing a corresponding taxonomy identifier (id). Since APRICOT has been designed to process multiple queries, the motif prediction can be carried out for the functional characterization of an entire proteome set corresponding to a taxonomy id. As the second input, users must provide a list of terms or keywords like names of domain families, Pfam ids or MeSH terms depending on the functional classes of interest, referred hereon as *domain selection keywords.* APRICOT uses a string-based search to select relevant entries from the domain resources, which are further utilized for identifying proteins that contain these domains. Optionally a set of terms called *result classification keywords* can be provided for the classification of predicted domains into smaller subsets in order to help users in navigating large datasets or classifying proteins by the functional similarity.

### 2. Modules for domain prediction and annotations

The core functionalities of APRICOT involve a multi-step process for the selection of proteins by identifying functional sites or domains of interest in their sequences followed by their annotations by various biological features. We have used a multifunctional human protein PTBP1 (47) as an example in order to describe the different modules involved in domain prediction and annotations in Figure 2. PTBP1 is an mRNA regulator that contains several repeated RBDs, specifically a highly abundant eukaryotic domain called RNA Recognition Motifs or RRMs (48).

**Fig. 2.**
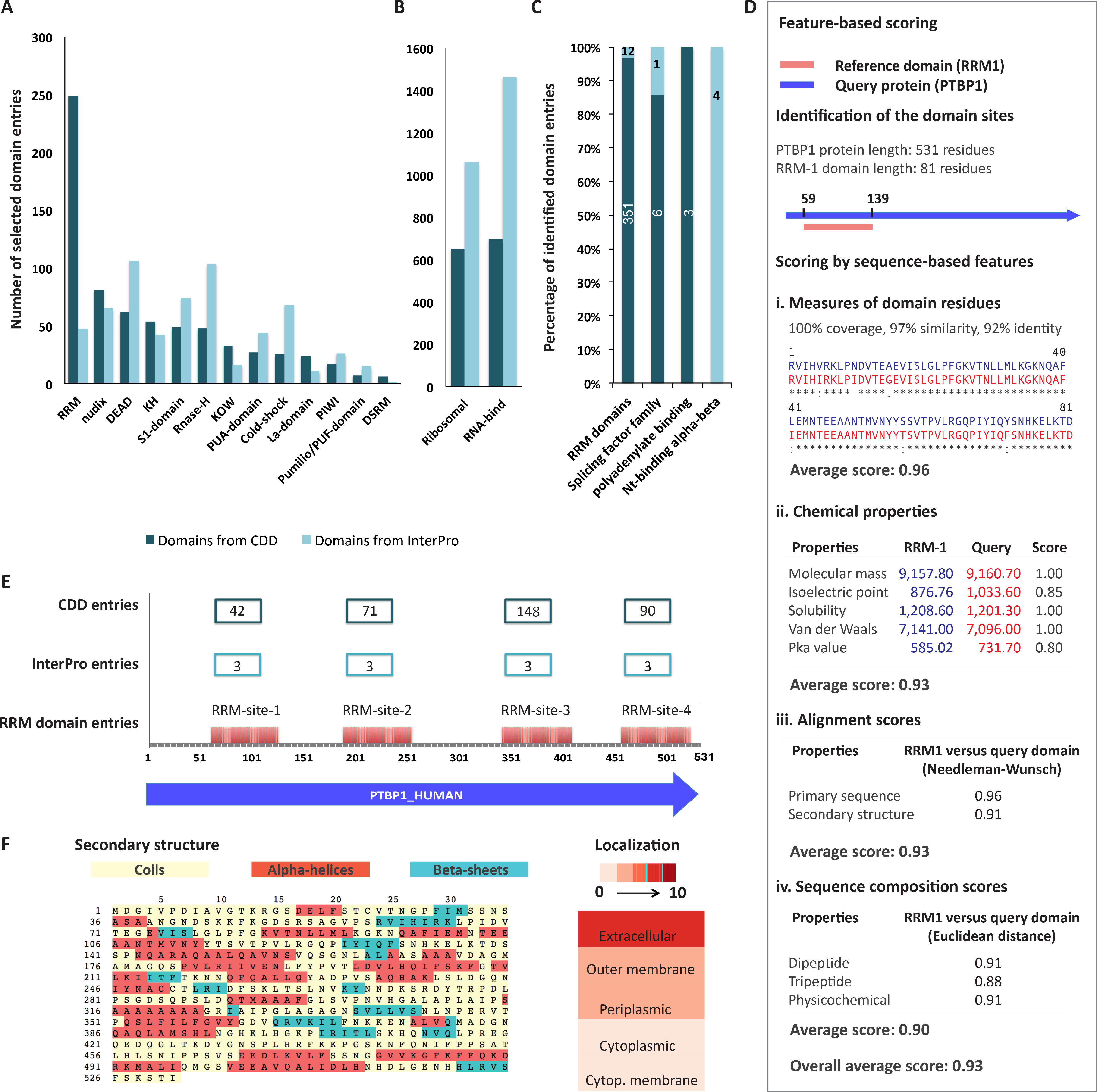
Different components of APRICOT for the characterization of RBPs are explained using an example of a human protein, PTBP1. (A) Bar chart showing the distribution of the known RNA binding domains collected from the CDD and the InterPro consortium. Several of these domains were selected by more than one domain selection term. (B) Additional domains selected by RNA-binding ribosomal domains and the term ‘RNA-bind'. (C) Domain entries from CDD and InterPro database, which were identified by APRICOT in the PTBP1 human protein. (D) A schematic workflow illustrating different processes involved in feature-based scoring resulted from a comparative analysis of RRM-1 domain (RRM1_PTBP1) and the corresponding domain identified in PTBP1 human protein. As shown in the schema, the features involved in this analysis have been classified into four categories, each comprising of specific set of sequence-based features. The features are scored by Bayesian probabilities in a range of 0 to 1, where 1 signifies a complete match between the reference and the domain identified in the query. (E) The 4 RRM sites in PTBP1 protein corresponding to different RRM entries from CDD and InterPro. (F) Visualization of additional annotations of PTBP1 protein by secondary structure and probability of sub cellular localizations generated by APRICOT.

2.1 Selection of reference domain set: A string based selection of domain families and functional motifs are carried out using the *domain selection keywords* to create a reference domain set. From the collections of domain entries from different domain databases in the CDD and InterPro consortiums, domains are selected when they match at least one of the provided terms in their annotations. In this analysis, we considered the domains obtained from the human interactome study (1, 4) as the comprehensive resources for building a reference RBD set. To report high confidence RBPs by avoiding the selection of ambiguous and functionally irrelevant domains, we included all the classical RBDs in *domain selection keywords* (Figure 2A). In order to account for ribosomal proteins, 109 terms related to RNA-binding ribosomal domains (4) were included in *domain selection keywords.* An additional term ‘RNA-bind’ was introduced to include any additional RBDs in the reference set that are well described as RBDs in databases but are not classified under classical RBDs (Figure 2B). Using these domain selection keywords, a total of 4,951 RBD entries were curated from CDD (1,951 entries) and InterPro (3,000 entries), referred as *reference domain set,* which was used for filtering domain predictions in the downstream analysis.

2.2 Domain prediction: In this step query amino acid sequences are characterized with all the possible domains from the databases without filtering a certain functional class. The sequences are subjected to domain prediction using RPS-BLAST and InterProScan to query their CDD and InterPro respectively (Figure 2C). By default, APRICOT uses both CDD and InterPro for the domain predictions, however users can choose one of the databases to reduce the run-time. Since the primary requirement of this module is the amino acid sequences of the query proteins in FASTA-format, users can analyse novel sequences even when the gene/protein ids are unknown or lacking.

2.3 Selection of proteins by functional domains of interest: This module allows the selection of relevant proteins from the query sets based on the predicted domains obtained in the previous step. The proteins are considered as candidates if they contain one of the domains of interest. Cut-offs for various statistical parameters (discussed below) can be defined for the selection of the predicted domains to identify such candidates, which are further annotated with additional information, such as ontology, pathway and cross-references to different databases.

2.4 Feature-based scoring: This module ranks the domain predictions by their relevance. For this purpose, a comparative analysis is carried out between the protein region that are predicted in the candidate proteins as domain of interest and the corresponding fragments of their reference consensus sequence. This comparison is done for a number of sequence-based features namely chemical properties (average mass, pKa and pI), alignment scores calculated by Needleman-Wunsch algorithm (primary sequence and secondary structure), Euclidean distance of protein compositions (di-peptides, tri-peptides and physico-chemical properties) and measure of homology between predicted sites and reference domains (for details see Supplementary Material S1A). A relative similarity between the predicted functional site and the reference domain consensus for these sets of features are calculated. We use Bayesian probabilistic score in a range from 0 to 1 to represent the functional potential of the predicted motifs, where 1 indicates the highest probability (Figure 2D). To further estimate the statistical significance of a predicted domain, P-values are calculated for the sequence-based features except for the chemical properties. These probabilistic scores and P-values allow users to select proteins with high confidence motif predictions.

2.5 Additional annotations of the selected proteins: Upon selection of proteins of functional relevance, users can choose to further annotate these proteins by information like sub-cellular localization by PSORTb (48), 8-state secondary structure by RaptorX (49), additional GO allocation and tertiary structure homologs (Figure 2E and Figure 2F further detail in the Supplementary Material S1B).

## 3. Program output

A comprehensive result is returned by APRICOT at each step of analysis and stored with relevant information that serves as the input for the subsequent steps. For example, the data for predicted domains can be repeatedly used for extracting proteins of different functional classes. The selected proteins are provided in a tabular format with the statistics on domain prediction and corresponding annotations obtained from UniProt and the comparative analysis (Supplementary Figure S2). To provide an easy navigation through the large-scale analysis data, the results can be classified using *result classification keywords,* into smaller subsets of proteins with enzymatic activities or specific functional aspect of proteins. Additionally, graphs and charts are provided to aid the visualization of the resulting data.

## Training sets

For the identification of the most suitable parameters and their corresponding cut-offs for domain selection, training sets were collected from the manually curated and reviewed subset of the UniProt Consortium-SwissProt (51). A positive set of proteins was selected by using the keyword ‘RNA-binding’. A second set of proteins was selected by using all the terms indicating functional association of proteins with nucleic acid. A third set comprising all the uncharacterized and hypothetical proteins from the database was selected. All these sets of proteins were subtracted from the SwissProt data and the remaining data consisting of 271,219 proteins were considered as the resource for negative set. All the redundant protein sequences from both positive and negative sets were removed by clustering the sequences using BLASTclust (52) using 90% of sequence identity. A total of 4,779 non-redundant proteins were compiled in the positive set and a set of 5,834 proteins were selected for negative set, referred to henceforth as SwissProt-positive and SwissProt-negative respectively (Supplementary Table S4).

## Test sets

To consistently evaluate the sensitivity, specificity and accuracy of APRICOT, a pair of positive and negative set was obtained from NCBI Reference Sequence (RefSeq), a non-redundant (nr) database (53), using the terms ‘RNA-bind’ and ‘periplasmic’ respectively. The former term retrieved 4,470 RBPs proteins from various organisms. The term ‘periplasmic’, which retrieved 5,836 bacterial periplasmic proteins, was considered as a resource for non-RNA-binding proteins based on the assumption that the majority of periplasmic proteins lack RBDs. Using BLASTclust from the NCBI-BLAST package (52) the proteins in each set were clustered by 75% sequence homology, which resulted into 687 proteins in positive set and 1,199 proteins in negative set, henceforth referred as nr-positive and nr-negative respectively. An additional pair of positive and negative set was obtained from RNApred webserver (11), which will be referred as RNApred-positive (377 proteins) and RNApred-negative (355 proteins). The sensitivity of the pipeline was tested on other positive datasets collected from various resources, which are RBPDB (53), RNAcompete (6), RBRIdent (18), rbp86 (55), rbp109 (55) and rbp107 (55) consisting of 1,101, 205, 281, 86, 109 and 107 proteins respectively. The datasets are listed in detail in the Supplementary Table S3.

In order to show practical applications of APRICOT as a tool for large-scale data analysis like complete proteome sets, two model organisms were evaluated on the genomic scale. *E. coli* K12 genome (taxonomy id: 83333) was used as an example for bacterial species and *Homo sapiens* (taxonomy id: 9606) was used as an example for eukaryotic species consisting of 4,479 and 70,076 protein entries in UniProt database respectively. The positive RBP sets were selected from both the proteomes to quantify the accuracy with which APRICOT identifies RBPs in these genomes. We considered 1,535 non-redundant human proteins as positive set (Supplementary Table S6), which were proposed as RBPs in the global experiment-based studies or were reported by independent publications (1-4). So far no global study has been reported for the genome wide identification of RNA-binding proteins in bacteria. Beside ribosomal proteins, only a few proteins such as Hfq (20), CsrA (21), YhbY (56), SmpB (57), ProQ (58), CspA (59) and CspB (59) have been reported as RBPs in *E. coli.* Hence, a larger RBP reference of *E. coli* K12 was retrieved from UniProt database using GO term GO:0003723) for RNA-Binding that comprised of 160 proteins including the known RBPs (Supplementary Table S7).

## Assessment criteria

The statistical parameters for domain predictions in the training set and the performance of the tool on the test sets were evaluated by using standard binary criteria of sensitivity (SN), specificity (SP), accuracy (ACC), Matthews Correlation Coefficient (MCC) and F-measure, using the following equations where TP, FN, TN and FP are true positive, false negative, true negative and false positive respectively.
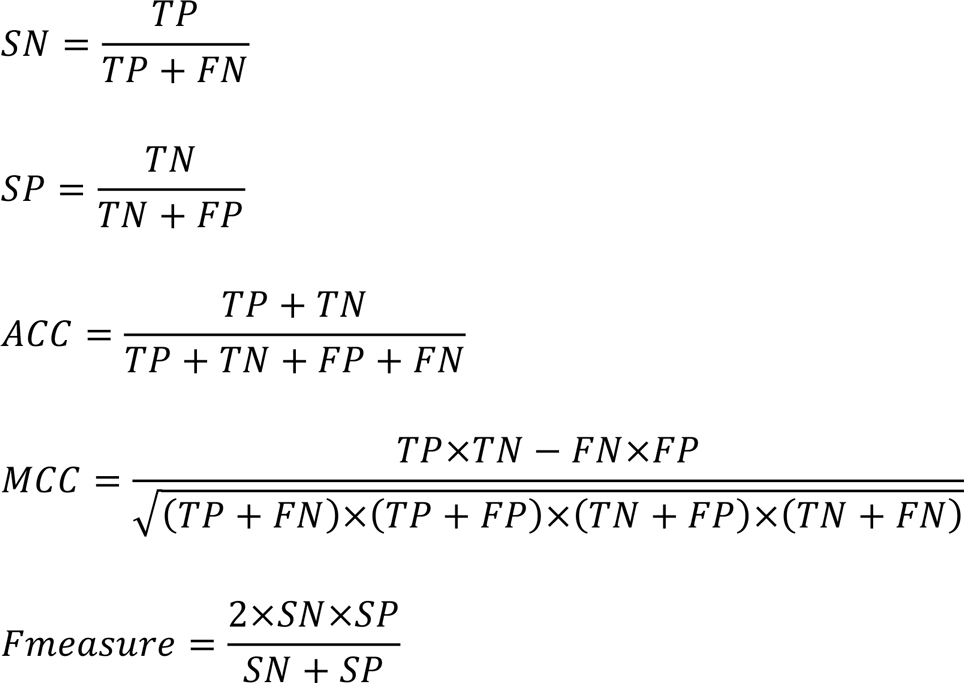

Receiver operating characteristic (ROC) curve and their area under the curve (AUC) was used as a criterion for accuracy, which was plotted using false positive rate (FPR or 1-SP) and true positive rate (TPR or SN).

## RESULTS

### Parameter optimization for the selection of predicted domains

The training sets, SwissProt-positive (4,779 proteins) and SwissProt-negative (5,834 proteins), were analysed in order to evaluate the ability of the method to accurately differentiate RBPs from non-RBPs. For this evaluation, we used statistical parameters of sequence similarity, residue identity, residue gap and E-value of the domain prediction to describe the similarity between a query and its corresponding reference. Unlike residue identity, sequence similarity accounts for the edit operations like positive substitutions, thereby capturing the secondary structure information at a better resolution. An E-value for searches of homologs against a database represents the likelihood that a given match in a sequence is purely by chance, meaning that a lower E-value reflects a higher significance of database match. We describe an additional parameter namely the domain coverage, which is the percentage of the length predicted as domain in the query compared to the original length of reference domain.Generally, lower domain coverage suggests a random homology of the predicted domain, whereas higher domain coverage reflects a higher potential of a domain to be functionally relevant.

Initially we investigated the analysis of the training sets by naïve approach, which involved InterProScan and CDD based batch-search methods in their default settings. Analysis by InterProScan achieved a TPR of 0.77 and CDD achieved a TPR of 0.79. Several queries in CDD based method were annotated as RBD containing proteins with coverage lower than 10% and sequence similarity lower than 5%, which indicated poor conservation of the functional domains. Similarly, InterProScan failed to characterize several RBPs due to its stringent filtering criteria. Interestingly, several RBPs were reported by only one of the methods, hence when the results from both the analyses were combined, an increased TPR of 0.82 was achieved. This clearly showed the potential to achieve higher sensitivity by the combined approach, which is implemented in APRICOT. We further analysed the training datasets by APRICOT,which predicted thousands of RBD entries in both positive and negative sets that were evaluated using systematically varying cut-offs of each parameter to optimize the identification of RBPs. The corresponding ROC curves were generated and optimal cut-off ranges were defined by identifying the values of the parameters that show a optimal TPR (closer to 1) and FPR (closer to 0) with high ACC (closer to 1), resulting into statistically significant AUC, MCC and F-measure (Figure3A and Supplementary Table S4).

For the coverage of the predicted domains, the minimum cut-off was recorded to be 39% that attained an accuracy, TPR, FPR, MCC value, and F-measure of 0.81, 0.87, 0.24, 0.63 and 0.81 respectively. Using a higher cut-off of 60% a lower TPR 0.81 but a better FPR 0.16 was obtained, which consequently shows better ACC and F-measure. Similarly, the optimal threshold for the minimum cut-off of sequence similarity was recorded to be 24%, which attains accuracy, TPR, FPR, MCC value of and F-measure of 0.81, 0.83, 0.20, 0.63 and 0.81 respectively. Similarly, as shown in the ROC curve, by using a minimum cut-off of 15% for the residue identity and at a maximum E-value cut-off of 0.01, high accuracies of 0.81 and 0.82 were achieved. The decision values of the parameters were further ranked, individually and in combinations, for all the predicted RBD entries in the training sets, we generated ROC curves and AUCs to identify their marginal contributions on overall accuracy in detecting RBDs (Supplementary Figure S5).

This evaluation led to the selection of domain coverage and sequence similarity as the default parameters for the APRICOT analysis with their minimum cut-offs of 39% and 24% respectively. The analysis by APRICOT using the selected parameters with their defined cut-offs achieves a TPR of 0. 85, which is higher than the naïve approach. The MCC and F-measure achieved for the APRICOT analysis of the training sets are 0.64 and 0.82 respectively. This successfully demonstrates the efficiency of the selected parameters and their cut-offs in identifying RBPs with high accuracy of 0.82.

### Assessment of the pipeline performance

A variety of positive datasets were analysed by APRICOT, on which the pipeline achieved sensitivity in a range of 0.81 to 1 (Figure 3B) demonstrating its high efficiency in domain-based characterization of RBPs. A more detailed evaluation of the pipeline performance was carried out on the paired dataset of nr-positive and nr-negative, and RNApred-positive and RNApred-negative (Table 1).

**Fig. 3.**
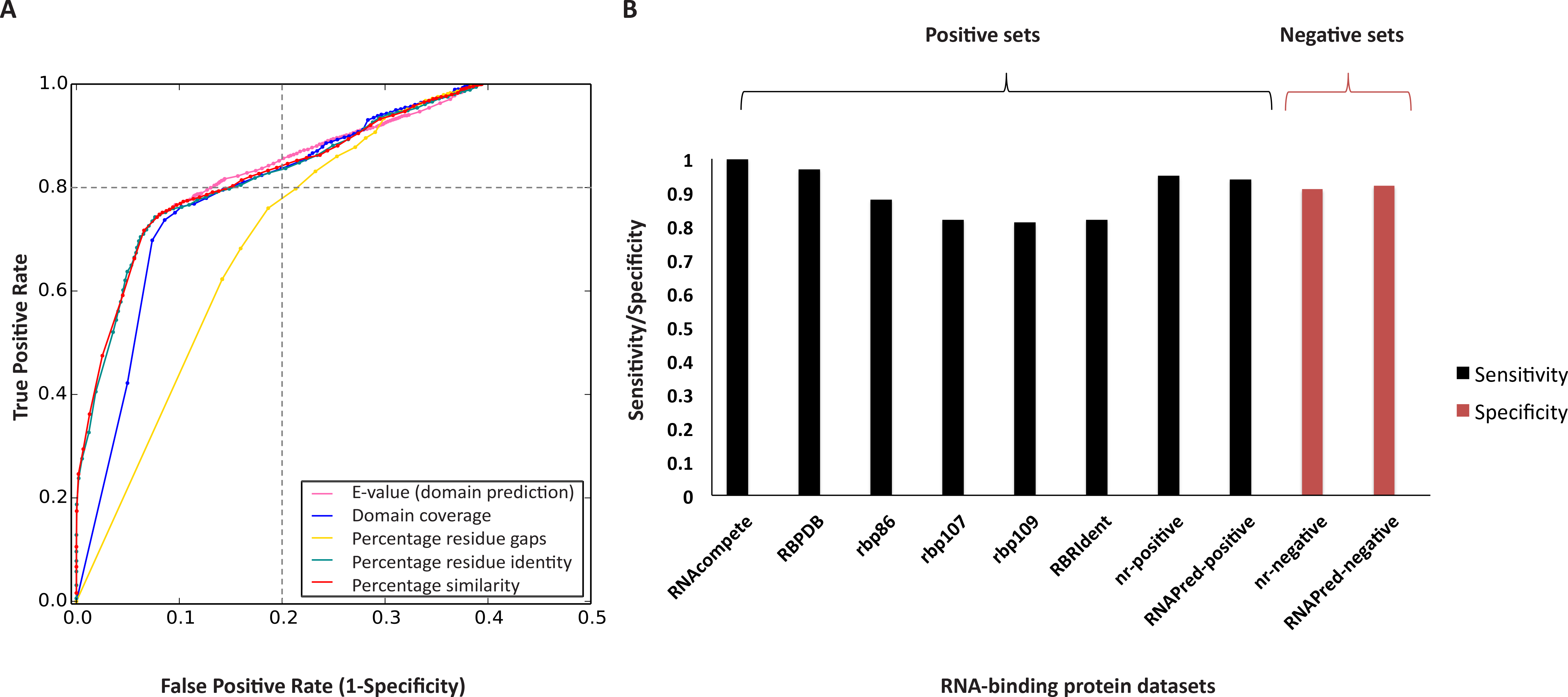
Selection of parameter cut-offs for RBP selection and the performance assessment ofm APRICOT on different datasets. (A) The ROC curves were generated for the domain prediction parameters of domain E-value (magenta), coverage (blue), residue gap (yellow), residue identity (green) and similarity (red). The optimal ranges for the parameters were defined for the selection of predicted domains at a considerably high accuracy (> 0.8 as indicated by the dashed lines) on the training sets (SwissProt-positive and SwissProt-negative). The minimum cut-off for most contributing parameters, percentage domain coverage and percentage similarity were recorded to be 39% and 24% respectively, which together attained an accuracy of 0.82. (B) The bar chart illustrates the performance of APRICOT on different datasets by means of sensitivity (shown in black) and specificity (shown in red). APRICOT was evaluated on 8 positive datasets and 2 negative datasets, which showed an average sensitivity of 0.90 and an average specificity of 0.91.

**Table 1.** Performance of APRICOT on positive and negative pair of datasets obtained from NCBI database and RNApred method.

| Datasets | RNApred | NCBI (nr) |
| --- | --- | --- |
| **Dataset types** |  |  |
| Positive set | 376 proteins | 687 proteins |
| Negative set | 355 proteins | 1,199 proteins |
| **Measures of performance assessment** |  |  |
| TP proteins | 344 proteins | 657 proteins |
| FP proteins | 47 proteins | 119 proteins |
| TPR (SN) | 0.96 | 0.97 |
| FPR (1-SP) | 0.10 | 0.13 |
| Accuracy | 0.93 | 0.92 |
| MCC | 0.86 | 0.85 |
| F measure | 0.93 | 0.92 |

To demonstrate the efficiency of APRICOT on large-scale data, the complete proteomes of *Homo sapiens* and *E. coli* K–12 were analysed. The human proteome set containing 70,076 UniProt protein entries was subjected to domain prediction. A known set of 1,535 non-redundant RBPs was used as positive reference set (4) of which 25 RBPs have not been defined with any RBDs. The reference domain set was considered for the initial identification of RBPs using pre-defined cut-offs for the aforementioned default parameters. Upon filtering of proteins by predicted domains, 1,091 from the reference RBP set were reported with at least one RBD from the reference domain set, showing a sensitivity of 0.71. By including the non-classical RBDs in the reference domain set, 68 more proteins could be recognized as RBPs and 201 RBPs could be recognized additionally by further including domains listed as RBDs unknown (Supplementary Table S6). The remaining 180 proteins that are not identified as RBPs by APRICOT do not contain RBDs and are listed as RNA-related proteins by Gerstberger *et al.* (4). The data for this analysis has been provided in the Supplementary Table S6.

A similar analysis of the complete proteome of *E. coli* K–12 was carried out by APRICOT using the default parameters with the reference domain set (Figure 2A and Figure 2B). In the initial characterization of RBPs, 673 sequences were selected as RBP candidates by RPS-BLAST and 502 sequences by InterProScan analysis. These proteins account for 806 RBP candidates, of which 369 proteins were identified as putative RBPs by both the methods. From the full proteome set, APRICOT could successfully identify all the known *E. coli* RBPs. Specifically, Hfq, CsrA, YhbY, SmpB, ProQ, CspA and CspB were identified due to highly conserved RBDs in their sequences, which have been previously reported and characterized for their regulatory roles (Table 2). Furthermore, from the GO term derived 160 RBPs from *E. coli* K–12, 129 were identified correctly by APRICOT that demonstrated a sensitivity of 0.80. APRICOT failed to identify the remaining 24 proteins as RBPs because either the predicted RBDs could not pass the parameter filters or the *reference domain set* lack specific domains associated with these proteins. These unidentified RBPs included CRISPR system Cascade subunits, toxic proteins and several enzymatic proteins like ribonucleases, tRNA-dihydrouridylases and mRNA interferases.

**Table 2.**
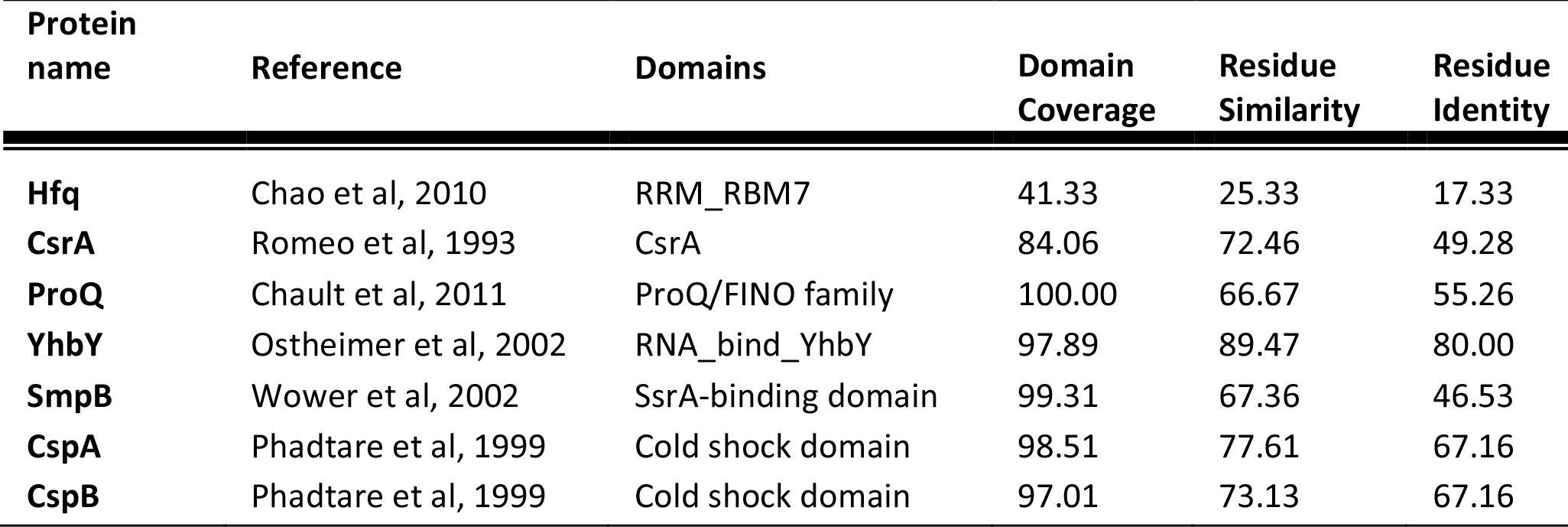
List of known RBPs in E. coli. These RBPs consist of well-defined RNA-binding domains. Hence, as shown in the table they were predicted to have higher coverage and similarity when compared with their reference domains.

| Protein name | Reference | Domains | Domain Coverage | Residue Similarity | Residue Identity |
| --- | --- | --- | --- | --- | --- |
| Hfq | Chao et al, 2010 | RRM_RBM7 | 41.33 | 25.33 | 17.33 |
| CsrA | Romeo et al, 1993 | CsrA | 84.06 | 72.46 | 49.28 |
| ProQ | Chault et al, 2011 | ProQ/FINO family | 100.00 | 66.67 | 55.26 |
| YhbY | Ostheimer et al, 2002 | RNA_bind_YhbY | 97.89 | 89.47 | 80.00 |
| SmpB | Wower et al, 2002 | SsrA-binding domain | 99.31 | 67.36 | 46.53 |
| CspA | Phadtare et al, 1999 | Cold shock domain | 98.51 | 77.61 | 67.16 |
| CspB | Phadtare et al, 1999 | Cold shock domain | 97.01 | 73.13 | 67.16 |

The feature-based scores were calculated for each domain selected from the predicted data,which facilitate in differentiating highly reliable RBD predictions from the low confidence RBD predictions. Query proteins that consist if high confidence RBDs were further annotated with additional information, namely subcellular localization, secondary structures, GO terms and tertiary structures (Supplementary Table S7).

These proteome-wide analyses clearly demonstrate a high sensitivity of the pipeline in identifying RBPs based on functional domains. However, it also shows a limitation related to the dependence of query characterization on the functional domains and motifs selected from the databases based on the user-provided terms.

### Identification of other functional classes by APRICOT

Importantly, in addition to the application for the functional identification of RBPs, APRICOT modules can be easily adapted for one or multiple other functional classes. As a part of the Critical Assessment of Function Annotation (CAFA), a project to assess the methods for computational annotation of protein functions (60), APRICOT was successfully used to annotate bacterial datasets comprising of more than 1 million proteins by a wide number of biological functions (arXiv:1601.00891 [q-bio.QM]). In order to emphasize the aspect of APRICOT as a tool for the characterization of other functional classes of proteins, we chose kinase proteins from *E. coli* (strain K–12) as the reference set. Kinases are known to catalyse the transfer of phosphate groups to a substrate molecule using ATP as a phosphate donor. In UniProt database, 110 proteins from *E. coli* (strain K–12) are annotated with various kinase activities (for e.g. Serine/threonine-protein kinase, Signal histidine kinase and Shikimate kinase) and are tagged by the GO term (GO:0016301) for kinase activity.

The APRICOT pipeline was supplied with the term ‘kinase', for the selection of reference domain set and the pipeline was applied to the kinase proteins (Supplementary Table S8). Out of 110, 106 kinase proteins were identified correctly by APRICOT, achieving a sensitivity of 0.96. The set of proteins that was not selected by APRICOT, contain kinase-associated domains that were not present in the reference domain set due to the pipeline domain selection constraints. This analysis suggests that APRICOT is efficient in the characterization of proteins based on pre-defined set of domains associated with functional classes other than RBPs as well. However, it should be noted that the accuracy of the results depends on the choice of terms for the domain selection.

### Comparative assessment of APRICOT with other RBP prediction tools

Although there are several approaches developed for the prediction of nucleic acid binding sites, we could compile only four tools described for their original aim to predict RBPs, namely SVMprot (61), RNApred (12), SPOT-Seq-RNA (63) and catRAPID signature (13). SVMprot was designed to predict RBPs by SVM based classification of proteins primary sequences into functional families (54 Pfam families) and it was made available as a webserver. Since the tool is no longer available, we could not include it in our comparative analysis. RNApred uses SVM models that are developed with amino-acid compositions and PSSMs. SPOT-Seq-RNA, uses structure homology based predictions of the RBPs and also allows the identification of the binding residues and binding affinities using SPARKS X (10) and DRNA tools (64) respectively. The fourth tool, catRAPID signature, is a SVM based method to identify RNA-binding proteins and their binding regions based on physico-chemical properties. We conducted comparative assessment of APRICOT's capabilities with these tools (Table 3).

**Table 3.** Comparative evaluation of APRICOT with other RBP prediction tools: catRAPID signature, SPOT-Seq-RNA and RNApred. Different features of the tools are listed in the upper part of the table to highlight the main differences and advantages over other tools. The comparative performance of the tools, shown in the lower part of the table, was assessed using the positive set of RBscore_R246 containing 246 proteins (total proteins from RBscore_R130 and RBscore_R116), and the negative set of RNApred-negative containing 355 proteins. APRICOT achieved higher overall sensitivity of 0.79 compared to the other tools.

| RBP prediction tools | APRICOT | catRAPID signature | SPOT-RNA-Seq | RNApred |
| --- | --- | --- | --- | --- |
| **Main features of the tools** |  |  |  |  |
| Main criteria for RBP characterization | RNA-binding motifs and domain families | Physico-chemical properties | RBP structure homologs | SVM classification by composition features of proteins |
| Additional analysis | Sequence-based scoring of domain features | Prediction of RNA-binding regions | RNA-binding residue prediction and binding affinity | PSSM based evolutionary information |
| Availability | Command-line | Webserver | Webserver and command-line | Webserver |
| Query types | Amino-acid sequences / gene names / UniProt protein / taxonomy ids | Amino-acid sequences | Amino-acid sequence | Amino-acid sequences |
| Allowed number of query proteins | Unlimited | 100 proteins or total number of submitted characters = 100000 | One query at a time | Unlimited for composition based or one query at a time for the PSSM based analysis |
| Probability scores for RBPs | Bayesian score (0-1) | SVM score (Threshold -0.2) | Z-score | SVM score (Threshold - 0.2) |
| Main criteria for RBP characterization | RNA-binding motifs and domain families | Physico-chemical properties | RBP structure homologs | SVM classification by composition features of proteins |
| **Performance assessment** |  |  |  |  |
| **TP (proteins)** | 193 | 125 | 166 | 180 |
| **FP (proteins)** | 44 | 150 | 6 | 102 |
| **TPR (SN)** | 0.79 | 0.51 | 0.67 | 0.73 |
| **FPR (1-SP)** | 0.12 | 0.42 | 0.02 | 0.29 |
| **ACC** | 0.83 | 0.54 | 0.83 | 0.72 |
| **MCC** | 0.66 | 0.1 | 0.69 | 0.44 |
| **F-measure** | 0.83 | 0.54 | 0.8 | 0.72 |

Unlike other tools, which have been trained or constructed on a certain set of reference set, APRICOT is established independent of any fixed set of reference because it selects reference domains for each analysis based on the user provided keywords. Therefore, it is capable of using any new RNA-binding domains that might be added in the integrated domain sources in future. APRICOT takes proteins that are predicted with statistically significant RBDs and scores them in comparison with their reference consensus sequence for various features using Needleman-Wunsch alignment scores, Euclidean distance and homology-based scores. At the end, the scores for each property are combined to obtain a Bayesian probabilistic score in a range of 0 to 1, where 1 indicates the best hits. The results from all the intermediate steps are provided to allow users to evaluate different statistical aspects of their study.

For an unbiased evaluation of the relative performances of APRICOT with RNApred, SPOT-Seq-RNA and catRAPID signature, we used two datasets RBscore_R130 (130 RBPs) and RBscore_R116 (116 RBPs), which are the training and test sets created for the RBscore_SVM approach in NBench (17). On RBscore_R130, APRICOT achieved a TPR of 0.88, whereas RNApred, SPOT-Seq-RNA and catRAPID signature attained much lower TPRs of 0.79, 0.82 and 0.55 respectively. On the RBscore_R116, which is indicated as a challenging set in NBench, APRICOT achieved a comparatively low TPR of 0.67, however, this was still higher than the TPRs achieved by RNApred (0.66), SPOT-Seq-RNA (0.51) and catRAPID signature (0.47). We also checked the performances of naïve RPS-BLAST, which is used for the batch-search of domain in CDD, and InterProScan, which is used for motif prediction in InterPro consortium. On both the datasets the naïve approaches for domain identification showed lower performances compared to their combined performance. Both the methods in their default setting achieved a TPR of 0.82 on the RBscore_R130 by identifying 107 RBPs. On the RBscore_R116, RPS-BLAST and InterProScan showed performances higher than SPOT-Seq-RNA but lower than APRICOT and RNApred by achieving TPR of 0.55 and 0.57 respectively.

APRICOT performed better than the other tested tools in all the assessment metrics used for the evaluation of RBscore_R246 (RBPs from both the datasets) as positive set and RNApred-negative (355 proteins) by achieving highest accuracy, MCC and *F*-measure of 0.88, 0.75 and 0.86 respectively (Table 3).

### APRICOT versus tools for the prediction of RNA-binding residues

A comparative assessment of the programs developed for the prediction of nucleic acid binding sites was carried out in NA Binding prediction Benchmark (17). Total 16 tools for the prediction of RNA-binding residues, 5 tools for the prediction of DNA-binding residues along with several datasets obtained from the structures of protein-nucleic acid complexes were included in this study (available at http://ahsoka.u-strasbg.fr/nbench/index.html). The motivation behind developing APRICOT is noticeably different from the tools involved in NBench. APRICOT identifies RBPs among large-scale query sets and further characterizes them by biological functions, whereas the 16 tools in NBench predict RNA-binding residues in the pre-defined RBPs. Practically, APRICOT and these tools can complement each other by first using APRICOT to identify RBPs and their corresponding RBDs and then applying the best performing NBench tools to obtain a high-resolution annotation by identifying RNA-binding residues. To evaluate the potential of this idea, we acquired 3,657 PDB entries, consisting of 24 different RNA related datasets in NBench selected at a resolution cut-off of 3.5 Å. This dataset was subjected to analysis by APRICOT and a comparative assessment was carried out between the identified RBD sites and the nucleic acid binding residues at the distance cut-off of 3.5 Å in each PDB entry (Supplementary Table S9).

We observed that the RNA-binding residues of 3,340 (91%) PDB entries overlap with the APRICOT predicted RBD sites showing an overall sensitivity of 0.91 (Figure 4A and 4B). The NBench tools were ranked by their sensitivities to identify RNA-binding residues together with APRICOT for its ability to identify RNA-binding sites on 24 datasets. As shown in Figure 4C, APRICOT was among the best performing tools compared to the other tools in NBench across the 21 diverse datasets. In agreement with the observations made for the tools, APRICOT showed a lower sensitivity on the New_R15 set (15 new structures) and RBscore_R116 (116 proteins, mentioned as difficult set). Furthermore, unlike most of the tools that do not show discriminative potential for RNA and DNA binding residues, APRICOT showed a high specificity (0.70) when 1,374 DNA binding proteins were included in the analysis. This evaluation demonstrates that APRICOT's domain prediction based analysis is an extremely efficient approach to identify RBPs and their corresponding potential RNA-binding region in the query sequences. Furthermore, it also implies that the resolution of the RBP studies could be enhanced significantly by first identifying the RBPs using APRICOT, followed by the analysis with the tools for the identification of RNA-binding residues in the predicted RBD sites.

**Fig. 4.**
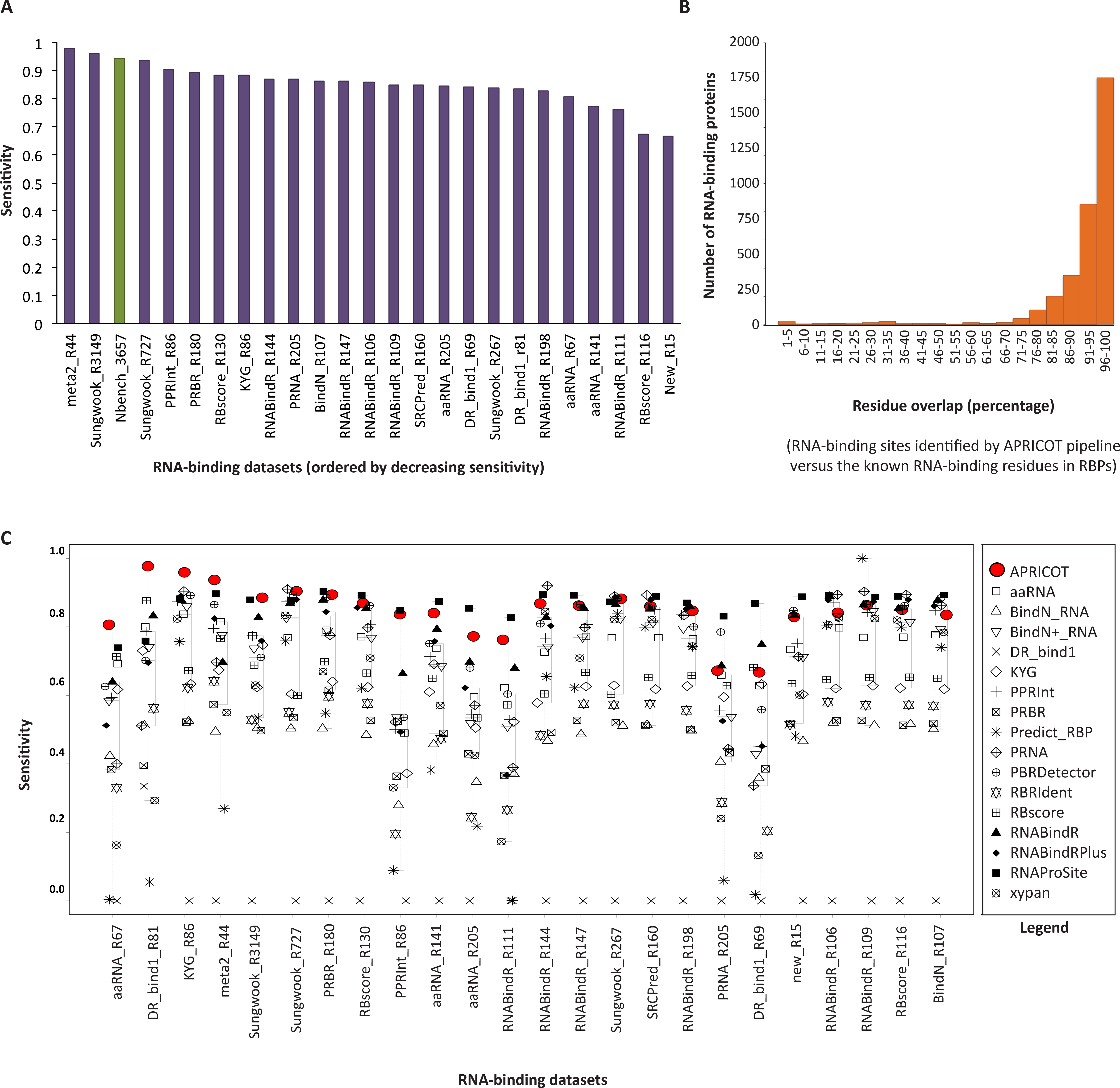
APRICOT based analysis of 24 datasets compiled from PDB in the NBench study for the evaluation of the tools for RNA-binding residue prediction (A) The bar chart showing the specificities achieved by APRICOT on different datasets, including the entire set of 3,657 RBPs (NBench_3657 shown in green). (B) Distribution of RNA-binding proteins based on the percentage of overlapping RNA-binding residues defined in NBench with the RNA-binding sites identified by APRICOT. The RNA-binding sites were identified in 3,445 of NBench_3657, of which 3,304 proteins have more than 70% of their RNA-binding residues overlapping with the RNA-binding sites. (C) Boxplots showing the sensitivities achieved by APRICOT in identifying RNA-binding sites (in red) and other RNA-binding residue prediction tools in identifying RNA-binding residues (in black) on NBench datasets. On all the datasets, APRICOT achieved sensitivities higher than or as good as high performing tools.

## CONCLUSIONS

APRICOT is an integrated pipeline for the sequence-based identification and annotation of the query proteins based on the functional motifs and domains of interest known from the experimental data. Notably, here we report APRICOT primarily as a tool for the sequence-based identification of RBPs, which uses a consistent set of reference RBDs derived from large-scale experimental studies. Using several domain data-resources and associated tools, the domains are predicted in the queries and only those proteins that contain domains of interest are further characterized. By involving a wide range of biological features for the characterization of functional motifs, the pipeline carries out an intensive comparative analysis between the predicted domains and their respective reference consensus. This comparison is translated into statistical scores that enable users to differentiate proteins that are predicted to harbour domains of high similarity with their reference sequences from proteins that have poorly conserved domains. The proteins are subjected to annotation by additional biological properties, such as subcellular localization and secondary structure to get further insight into their functional relevance.

The pipeline has been extensively tested on several RBPs and is optimized for the identification of RBPs in large datasets, such as complete proteomes of human and *E. coli.* For instance, APRICOT could successfully identify the respective motifs of CsrA, ProQ, YhbY and SmpB in *E.coli* with domain coverage higher than 80% and residue similarity closer to 70%. In addition to these previously characterized RBPs, APRICOT predicted a number *E.coli* proteins that can potentially interact with RNAs via RBDs, and hence, could be further validated by experimental studies.

A thorough comparison between APRICOT and the other RBP prediction tools successfully demonstrated its superior performance and efficiency in a wide range of datasets for the identification of RBPs. Furthermore, we showed that the RBD sites obtained from APRICOT analysis have high overlap with the known RNA-binding residue sites in RBPs. Hence, we suggest that analysis of APRICOT can be complemented with the RNA-binding residue prediction tools to achieve a high-resolution binding information of RBPs. Due to the automated framework and accessibility of different modules of the pipeline, APRICOT can be conveniently adapted for the characterization of other functional classes. In agreement, by applying the pipeline for the identification of kinase proteins in *E. coli,* we demonstrate that the tool is not built on a fixed set of domain information, but instead it allows users to characterize proteins based on the functional classes of their interest.

## AVAILABILITY

APRICOT is implemented in Python as a standalone command-line program, which can be executed on Unix systems. The tool has been extensively refined based on the requirements and suggestions by experimental researchers. The source-code for the command-line tool is available under the ISC license at https://pypi.python.org/pypi/bio-apricot(GitHub repository: http://malvikasharan.github.io/APRICOT/ and the releases are automatically submitted to zenodo for the current version 1.1).http://dx.doi.org/10.5281/zenodo.51917

## ACKNOWLEDGEMENT

The authors thank the members of the Vogel group, Eulalio group and the Core Unit (CU) Systems Medicine, especially Dr. Charlotte Michaux and Caroline Taouk for contributing into the project with their experimental knowledge and constructive feedback. The authors are grateful to Prof. Dr. Thomas Dandekar, Dr. Lars Barquist and Dr. Stanislaw Gorski for critically reading the manuscript and providing useful feedback.

## FUNDING

The Vogel and Eulalio groups are supported by funds from DFG and the Bavarian Ministry of Sciences, Research and the Arts in the framework of the Bavarian Research Network for Molecular Biosystems (BioSysNet).

## Authors Contributions

M.S. and J.V. designed the study. M.S. established the workflow, acquired the datasets, analysed and interpreted the results. K.U.F. provided useful technical supervision and contributed to the code development. A.E. and J.V. provided important feedback and discussions throughout the project. M.S. wrote the manuscript, which all co-authors extensively discussed and commented on.

## Conflict of Interest

None declared.

